## supplementary document for "Optogenetic spatial patterning of cooperation in yeast populations"

### Supplementary Information: Optogenetic spatial patterning of cooperation in yeast populations.

### Model of the Cooperator/Cheater system

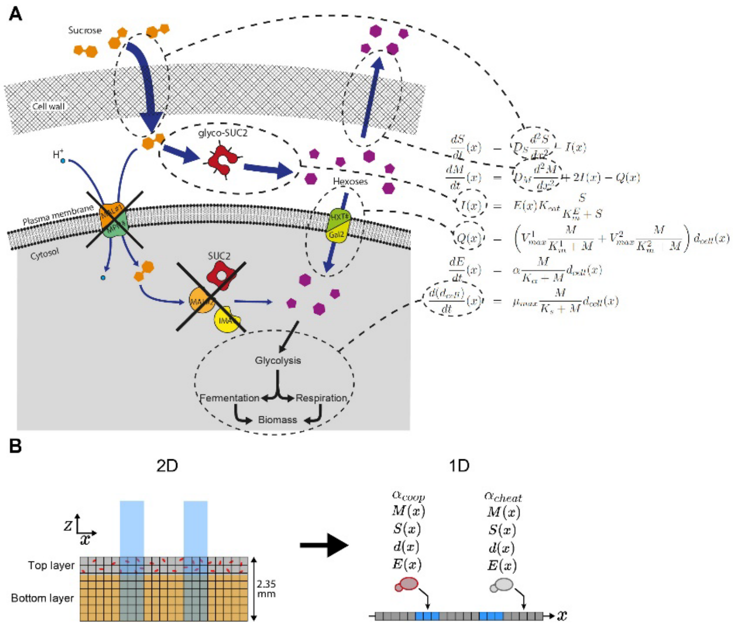

**Modelling yeast growth on sucrose**. (**A**) *Saccharomyces cerevisiae* sucrose pathway and the corresponding set of partial differential equations used to model the system. (**B**) Schematic view of the model (left) and its dimensional reduction to a one-dimensional problem (right). The geometry of the gel allows us to assume that diffusion in the *z*-axis happens very rapidly compared to diffusion in the *x* and *y* directions: ΔZ = 2.35 mm << ΔX = ΔY = 53 mm. We can thus simplify the model by considering the gel as homogeneous in the *z*-axis. We take dilution into account using the layer thicknesses (1.68 mm Phytagel and 0.67 mm agarose) to convert the sucrose concentration and cell density values.

To model the growth of yeast in sucrose, we use Michaelis–Menten kinetics for the enzymatic reactions (invertase hydrolysis Eq(3)) and high and low affinity glucose transporter Eq(4) and the Monod equation for the yeast growth rate Eq(6).

$$\frac{dS}{dt}(x) = D_S \frac{d^2S}{dx^2} - I(x) \tag{1}$$

$$\frac{dM}{dt}(x) = D_M \frac{d^2M}{dx^2} + 2I(x) - Q(x) \tag{2}$$

$$I(x) = E(x) K_{cat} \frac{S}{K_m^E + S} \tag{3}$$

$$Q(x) = \left( V_{max}^1 \frac{M}{K_m^1 + M} + V_{max}^2 \frac{M}{K_m^2 + M} \right) d_{cell}(x) \tag{4}$$

$$\frac{dE}{dt}(x) = \alpha \frac{M}{K_\alpha + M} d_{cell}(x) \tag{5}$$

$$\frac{d(d_{cell})}{dt}(x) = \mu_{max} \frac{M}{K_S + M} d_{cell}(x) \tag{6}$$

Details of the different variables and constants are summarized below:

| Variable | Description | Unit | Initial value | Source | Reference |
|---|---|---|---|---|---|
| M | Hexose concentration | M | 1.00E-08 | Fixed experimentally | |
| S | Sucrose concentration | M | 0.029 | Fixed experimentally | |
| E | Invertase concentration | M | 1.00E-35 | Fixed experimentally | |
| α | Enzyme production rate | mol.s$^{-1}$.cell$^{-1}$ | 1.8E-24 or 1.5E-25 | Fit | |

| | | | | | |
|---|---|---|---|---|---|
| d | Cell density | cell.$L^{-1}$ | 2.85E+09 | Fixed experimentally | |
| I | Invertase activity | M.$s^{-1}$ | - | Computed | |
| Q | Hexose consumption | M.$s^{-1}$ | - | Computed | |
| $D_S$ | Diffusion of sucrose | $m^2.s^{-1}$ | 6.10E-10 | Fixed experimentally | [1] |
| $D_M$ | Diffusion of hexoses | $m^2.s^{-1}$ | 7.60E-10 | Fixed experimentally | [1] |
| $K_{cat}$ | Kcat of monomeric Invertase | $s^{-1}$ | 4700 | Fixed experimentally | [2] |
| $\mu_{max}$ | Maximal growth rate | $s^{-1}$ | 7.50E-05 | Fit | |
| $V_{max1}$ | Maximal consumption rate | mol.$s^{-1}$.cell$^{-1}$ | 4.18E-17 | Fixed experimentally | [3] |
| $V_{max2}$ | Maximal consumption rate | mol.$s^{-1}$.cell$^{-1}$ | 2.60E-17 | Fixed experimentally | [3] |
| $K_{m1}$ | Affinity constant for hexoses | M | 0,0008 | Fixed experimentally | [3] |
| $K_{m2}$ | Affinity constant for hexoses | M | 0,021 | Fixed experimentally | [3] |
| $K_s$ | Monod constant | M | 0,00012 | Fixed experimentally | [4] |
| $K_m^E$ | Km of invertase | M | 0,026 | Fixed experimentally | [2] |
| $K_\alpha$ | Invertase production | M | 1.00E-05 | User defined | |

**Table –** Summary of the model parameters for yeast metabolism and growth on sucrose.

#### Parameter adjustment

We manually tuned these parameters so that the simulations fit both the dynamic and final densities of the corresponding experiments. We found the following values: μmax = 0.27 $h^{-1}$, $\alpha$coop = 1.8 $10^{-24}$ mol.$s^{-1}$.cell$^{-1}$ (corresponds to the maximal DMD light intensity) and $\alpha$cheat = 1.5 $10^{-25}$ mol.$s^{-1}$.cell$^{-1}$ (corresponds to the promoter leaking and the minimal DMD light intensity).

#### Simulation

To solve the PDEs, we use a Python package called *scikit-fdiff* (https://scikit-fdiff.readthedocs.io). We chose the Crack-Nicholson scheme to compute the diffusion of molecules across a discretized space and used reflective boundaries. We use a simulation hook to compute non-linear terms and prevent negatives values. We ran all simulations on a Intel(R) Xeon(R) CPU E5-1650v4 3.60 GHz processor and 64 Go of RAM.

### Building the OptoCube

Building an OptoCube is relatively straightforward. We used a static incubator with relatively large dimensions, a Digital micromirror device, a microcontroller, and a flatbed scanner. We propose here a set of devices that worked for us, but other commercial or DIY alternatives are of course possible. Details and scripts can be found here: https://github.com/Lab513/DIY_OptoCube. Briefly, the DMD was mounted on an incubator rack (OpenBeam construction kits). The scanner was placed below, facing up. The distance between the scanner glass and the DMD lens was 39 cm.

- Incubator: Memert (any sufficiently large incubator will do)
- DMD: DLP® LightCrafter™ 4500 TI
- Scanner: Canon LIDE400. Note that we opened the glass cover and applied black tape on some parts of the bottom of the scanner that are reflecting surfaces (this prevents undesired optogenetic activation due to reflected light artefacts).
- Microcontroller: Arduino UNO.

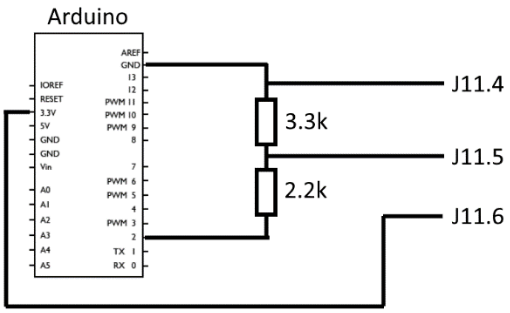

**Controlling the DMD with the Arduino board.** The DMD was powered with a 12 V 4.16 A power supply and connected to the Arduino by pins J11.4, J11.5, and J11.6 using a molex 51021-0600 connector. The scanner was connected to the computer with a USB cable. The Arduino UNO was also connected to the computer with a USB cable and used to control the DMD via pin number 2 to apply 3.3 V at the J11.5 pin through a voltage divider (2.2 kOhm and 3.3 kOhm). The Arduino 3.3 V pin was connected to the J11.4 pin and the ground pin, to the J11.6. We used Jupyter notebook to drive the OptoCube. The following packages were installed:

- serial (https://pypi.org/project/pyserial/)
- time (https://docs.python.org/fr/3/library/time.html)
- os (https://docs.python.org/fr/3/library/os.html).

The DLP® LightCrafter™ 4500 TI is a digital micromirror device (DMD) composed of 912 × 1140 micromirrors that can switch ON or OFF to reflect the light emitted from integrated LEDs (red, green, and blue). The projected intensity of each micromirror is controlled by pulse width modulation. One important technical limitation is that even when the mirrors are completely OFF, a significant amount of light leaks out of the DMD onto the projected surface. The DMD offers two main modes of projection: video mode or sequence mode. Video mode is performed through classical HDMI communication with a computer, allowing for straightforward and dynamic patterning. However, the range of light intensity in video mode is quite low, with high leakage. Thus, we used the sequence mode, which provides a better intensity range (from 0.0014 mW.cm$^{-2}$ to 1.13 mW/.cm$^{-2}$ for the blue LED). The main drawback of this mode is the lack of flexibility to change the mask being projected by the DMD. The mask must be loaded in the

DMD before the experiment starts and the maximum number of 8-bit masks that can be stored is six. We used the DLPLCR software ("DLPLCR4500EVM-GUI", which can be downloaded here https://www.ti.com/tool/DLPLCR4500EVM) to create and load the masks and to design the pattern sequence. More information on the procedure can be found in the DMD user guide. Briefly, the workflow is as follows:

1. Create an 8-bit image in BMP format of dimensions 912 x 1140 (width x height). The pattern must be drawn with 200% deformation in the vertical axis (due to the diamond shape of the mirrors; see the DLPLCR documentation for details)
2. This image is transformed by the DLPLCR software to a 24-bit BMP image.
3. Load the 24-bit image to the firmware using the DLPLCR software.
4. Connect the computer to the DMD using a USB cable and power the DMD.
5. Load the firmware in the DMD.
6. Edit and save the pattern time sequence in the DMD. You can then unplug the computer from the DMD.

**Setting up the VueScan© software to drive the scanner.** We used the following parameters:

- Media: "Black&White"
- Media size: A4
- Output file: 16-bit greyscale .tiff (no reduction nor compression)
- Make Grey from: Auto
- Scan resolution: 600 dpi
- Number of passes: 1
- Color balance: "None"

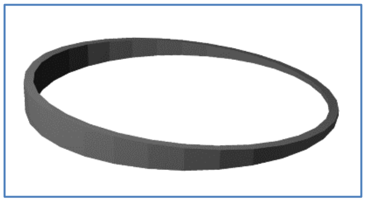

**Additional Requirements.** The Petri dish lids can generate imaging artefacts by reflecting light. To prevent this, we used a specially designed 3D printed part ("PlateTilterAngle5") to tilt the lids of the Petri dishes at 5°. Note that to maximize the image quality, you should ensure that the agar plates are:

1. Highly transparent to avoid light scattering (this is why we relied on Phytagel).
2. The gel layer is thin to allow the yeast layer to be in the focal plan of the scanner.
3. The lid should be coated with a surfactant solution to reduce droplet formation due to condensation. We used Triton 100X 0.05% (*v/v*) in 20% ethanol.

### Supplementary Table 1

| Name | HO locus | SUC2 locus | HIS locus | Nuclear marker | Background |
|---|---|---|---|---|---|
| yPH_428 | N/A | N/A | N/A | N/A | BY4741 |
| yPH_436 | N/A | N/A | EL222-HIS3 | N/A | BY4741 |
| yPH_449 | N/A | N/A | EL222-HIS3 | HTB2::mApple-Kan | BY4741 |
| yPH_457 | N/A | ΔSUC2 | EL222-HIS3 | HTB2::mApple-Kan | BY4741 |
| yPH_470 | pC120-Venus-SUC2 | ΔSUC2 | EL222-HIS3 | HTB2::mApple-Kan | BY4741 |
| yPH_471 | pC120-SUC2-P2A-Venus | ΔSUC2 | EL222-HIS3 | HTB2::mApple-Kan | BY4741 |
| yPH_484 | N/A | P2A-Venus | EL222-HIS3 | HTB2::mApple-Kan | BY4741 |
| yPH_536 | pC120-SUC2-HairPinC09 | ΔSUC2 | EL222-HIS3 | HTB2::mApple-Kan | BY4741 |
| yPH_540 | pC120GAL1-SUC2-HairPinC09 | ΔSUC2 | EL222-HIS3 | HTB2::mApple-Kan | BY4741 |

**Supplementary Table 1.** List of yeast strains used in this study.

### Supplementary Table 2

| Name | Use | Plasmid type | MoClo part type | Description |
|---|---|---|---|---|
| pPH297 | Repair fragment for homologous recombination (HR) with auxotrophic selection | x | x | EL222-HIS fragment by digestion |
| pPH330 | PCR template for repair fragment for HR with antibiotic selection | FA6a | x | HTB2-mApple-Kan PCR |
| pPH162 | Integration of optogenetically controlled genes | Expression cassette for CRIPSR/Cas9+gRNA | x | pML104-HO |
| pPH362 | SUC2 deletion | Expression cassette for CRIPSR/Cas9+gRNA | x | pML104-SUC2 |
| pPH374 | SUC2 tagging with -P2A-Venus | Expression cassette for CRIPSR/Cas9+gRNA | x | pML104-SUC2 |
| pPH296 | PCR template for pC120 integration in a “level 0” (lvl0) entry vector | x | x | pC120 PCR |
| pPH349 | PCR template for GAL1 minimal promoter to build p2RGal | x | x | pGAL1 PCR |
| pYTK97 | Golden Gate assembly to construct “level 1” (lvl 1) MoClo plasmid | MoClo lvl0 | 2 | pC120 |
| pYTK111 | Golden Gate assembly to construct lvl 1 MoClo plasmid | MoClo lvl0 | 3a | SUC2 |
| pYTK112 | Golden Gate assembly to construct lvl 1 MoClo plasmid | MoClo lvl0 | 3b | P2A |
| pYTK135 | Golden Gate assembly to construct lvl 1 MoClo plasmid | MoClo lvl0 | 3 | SUC2 |
| pYTK137 | Golden Gate assembly to construct lvl 1 MoClo plasmid | MoClo lvl0 | 2 | pC120GAL1_2R |
| pYTK140 | Golden Gate assembly to construct lvl 1 MoClo plasmid | MoClo lvl0 | 4a | HP: Hairpin mRNA degron C09 |
| pYTK118 | Repair fragment for HR with CRISPR | MoClo lvl1 | x | pC120-SUC2-P2A-Venus |
| pYTK119 | Repair fragment for HR with CRISPR | MoClo lvl1 | x | pC120-SUC2-Venus |
| pYTK147 | Repair fragment for HR with CRISPR | MoClo lvl1 | x | pC120-SUC2-HP |
| pYTK151 | Repair fragment for HR with CRISPR | MoClo lvl1 | x | p2RGal-SUC2-HP |

**Supplementary Table 2**. List of plasmids used in this study. We built several “level 0” plasmids compatible with the Modular Cloning framework[5] so we could easily assemble transcriptional units in “level 1” plasmids. All plasmids and sequences are available upon request.

### Supplementary Figure S1

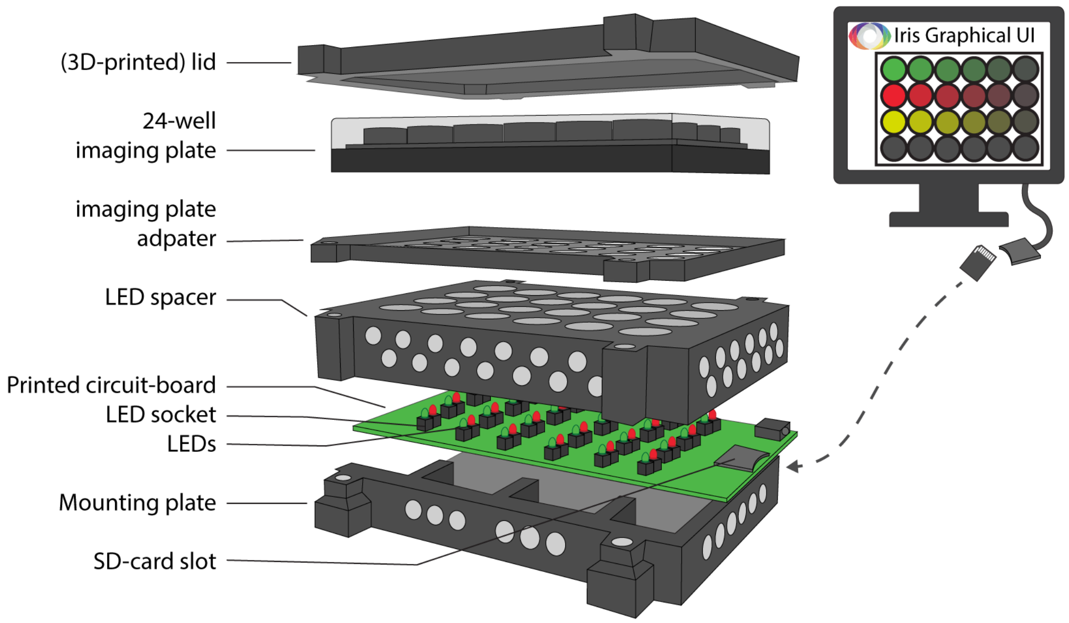

**Supplementary Figure S1**: 3D sketch of the Light Plate Apparatus[6] we used to activate SUC2 production with light. We routinely cultured yeast cells in 24-well plates and programmed the blue LEDs for each well to screen for optogenetic activation of SUC2. More details of the LPA can be found on the Tabor Lab's github (https://github.com/taborlab/LPA-hardware).

### Supplementary Figure S2

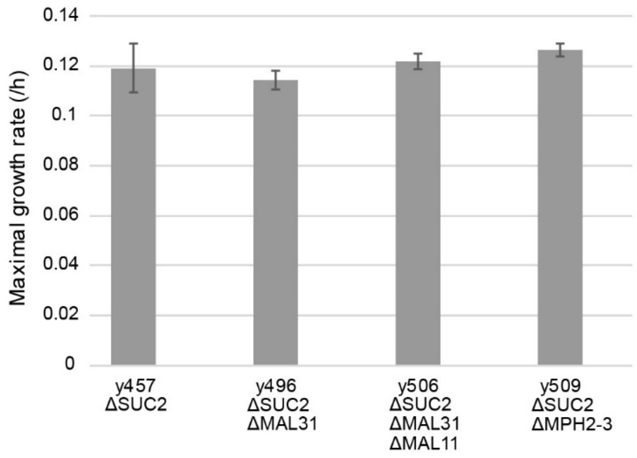

**Supplementary Figure S2:** Maximal growth rate of mutants of SUC2, MAL31, MAL11, and MPH2-3 in SC 1% sucrose obtained by measuring the increase in optical density over time.

### Supplementary Figure S3

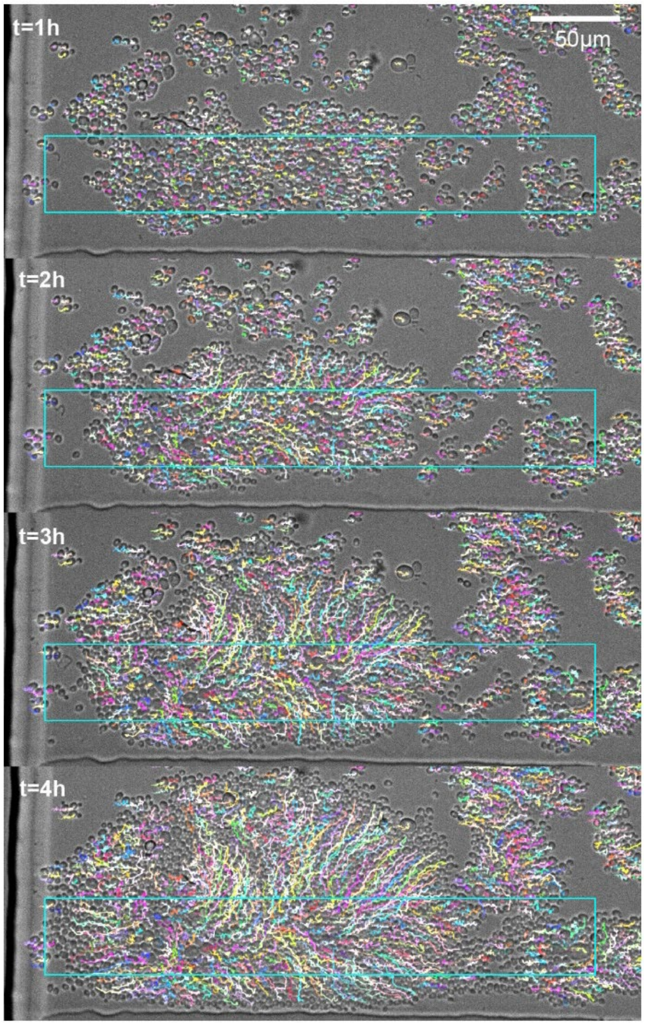

**Supplementary Figure S3:** Tracking of yeast cells (OptoSuc2) from Figure 3 (main text). Cells are growing in a microfluidic chamber perfused with 1% sucrose and a DMD was used to shine light on a given area (cyan rectangle). Tracking was performed with the TrackMate plugin[7] in ImageJ to confirm that growing cells push each other outside of the illumination area. Only high-quality trajectories starting at t = 0 h are plotted for better visibility. The (cyan) rectangle represents the area illuminated using the Mosaic (DMD) at 460 nm for 200 ms every 6 min.

### Supplementary Figure S4

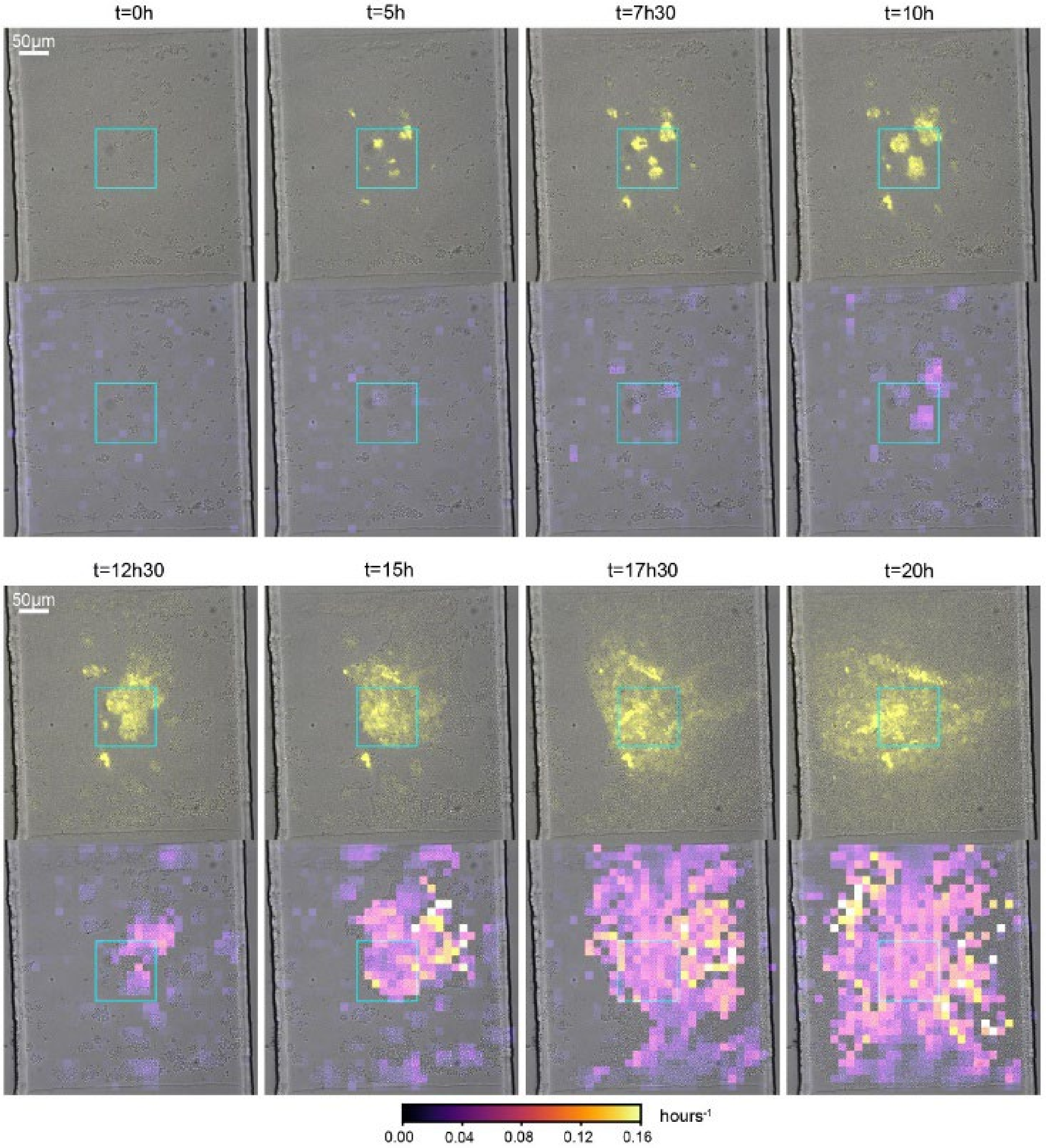

**Supplementary Figure S4:** Timelapse microscopy of SUC2-P2A-YFP strains (yPH_471) in a microfluidic chamber perfused with 1% sucrose. Cells within the cyan square were activated by light. We show the time series of an overlay of brightfield and yellow fluorescence images (top) and the divergence map of local cell velocities (bottom) to identify regions where cells are growing. The cyan square represents the area illuminated using the Mosaic (DMD) at 460 nm (intensity 20%) for 250 ms every 6 min. This experiment shows that cell growth leads to a flow of cells outside of the illuminated area. These cells are still marked by fluorescence because of the long lifetime of YFP. Similarly, we expect that these cells still express functioning Suc2p in their periplasmic space, which would allow the cells to process sucrose and to grow. As a result, growth is rapidly not constrained to the illuminated area but invades the full area of the microfluidic chamber. The long lifetime of Suc2p (and YFP) prevented us from limiting the growth of cells to the illuminated area at such a small scale.

### Supplementary Figure S5

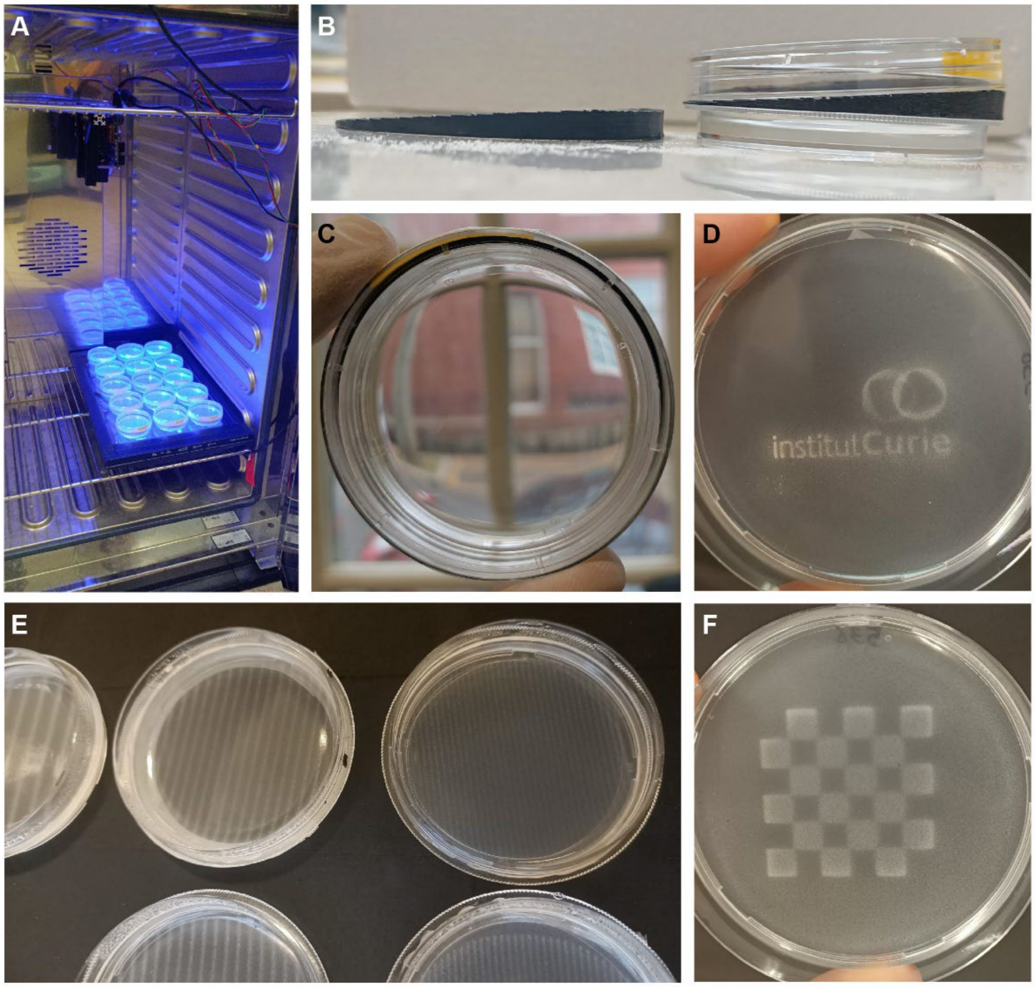

**Supplementary Figure S5.** Pictures of the OptoCube device and its application to activate growth at selected locations at the centimetre scale. (**A**) Interior of the OptoCube with the DMD (top) illuminating 15 plates placed on top of a scanner. The large incubator could accommodate a second DMD/scanner system, if required. (**B**) 3D printed part used to tilt the lids to 5°. This avoids rapid drying of the plates and avoids direct reflection of the DMD on the plates. (**C**) Image of an agar plate containing a two-layer gel with a homogeneous yeast suspension before light induction. The agar plate is transparent and exhibits homogenous cell density. (**D-F**) Examples of the growth patterns obtained when plates were illuminated with the Institut Curie logo (**D**), arrays of lines (**E**), and a chess board pattern (**F**).

### Supplementary Figure S6

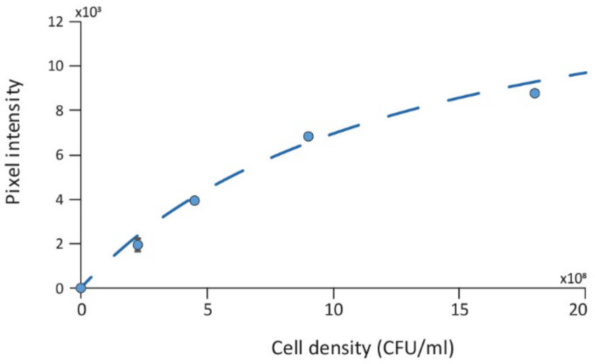

**Supplementary Figure S6**. **Calibration curve of the scanner** to determine the relationship between pixel intensity and cell density. We prepared agar plates with known cell densities in the top layer and measured the pixel intensities acquired by the scanner. We then fitted the data with the function $y = y_0 \frac{x}{x+x_0}$ with $y_0 = 1{,}6.10^4$ and $x_0 = 1{,}3.10^9$ and used this function to convert the pixel intensities into cell densities for our experiments. Round circles are data points, error bars represent standard deviation of duplicates, and the fit is plotted as a dashed line.

### Supplementary Figure S7

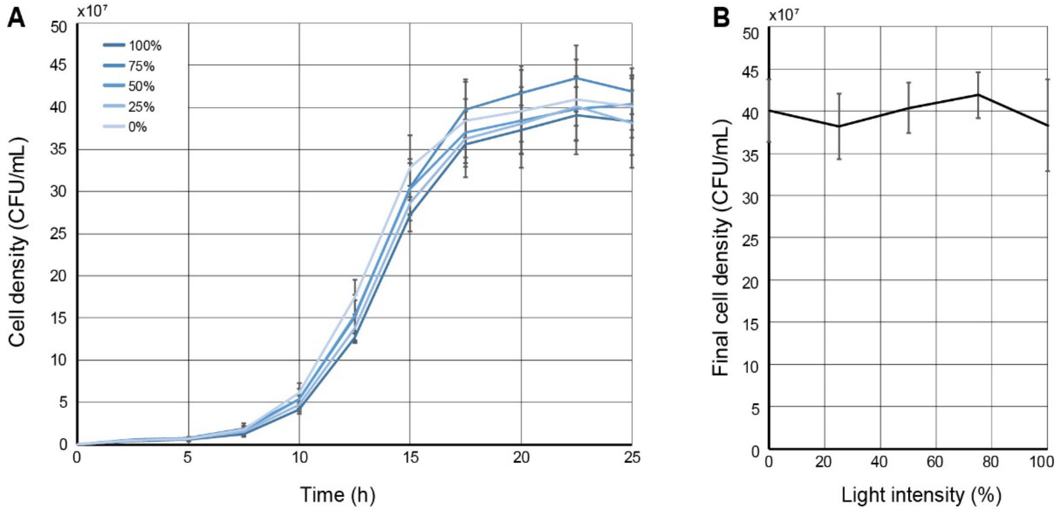

**Supplementary Figure S7**. **Growth of a homogeneous layer of cells**. We illuminated the entire surface of agar plates containing the WT strain and varied the light intensity from 0 to 100%. (**A**) Variation in the cell density as a function of time. Lines represent means of triplicate and error bars represent ± standard deviation. (**B**) Dependence of the final cell density on the light intensity at t = 25 h. No significant phototoxicity was observed in our system.

### Supplementary Figure S8

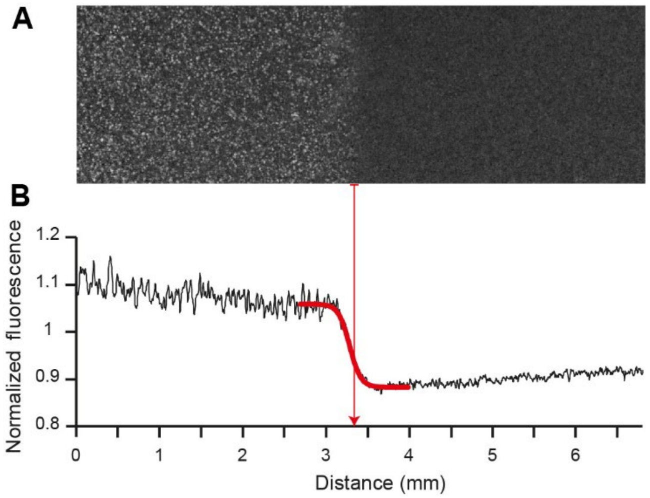

**Supplementary Figure S8.** Spatial resolution of the optogenetic induction of gene expression in the OptoCube. We applied a light pattern with a sharp transition from 100% to 0% of intensity for 24 h on a bilayered agar plate, with the top layer containing an optogenetic yeast strain that produces YFP under the control of the EL222/pC120 optogenetic system. (**A**) Normalized fluorescent image of the light pattern transition captured by a fluorescence macroscope and (**B**) quantification of the fluorescence intensity profile across the light-dark transition region. The normalization was performed by dividing the image by the image of a non-illuminated plate. The red line represents the fit of a sigmoid function $y=1/(1+\exp(-\lambda x))$ with parameter $\lambda$ = 14.95 $mm^{-1}$. As shown in the figure, the transition is sharp and typically of the order of half a millimeter.

### Supplementary Figure S9

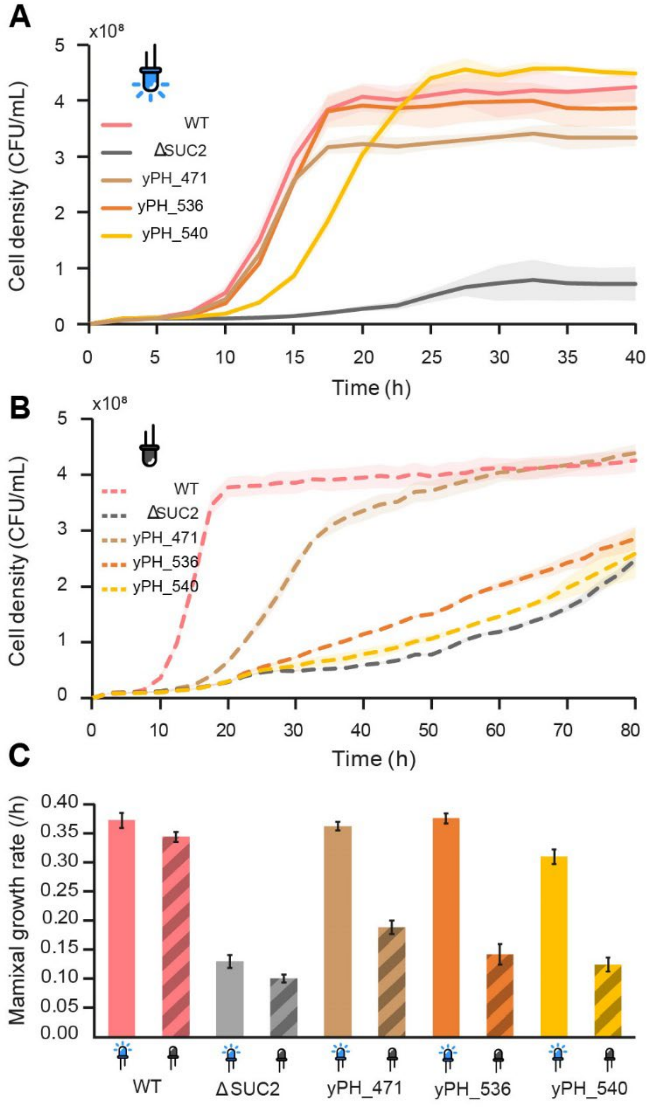

**Supplementary Figure S9. Growth curves of different optogenetic strains with homogeneous illumination in the OptoCube obtained in 1% sucrose. (A, B)** Growth in the dark (control) **(A)** and 100% light intensity (**B**). Lines represent the mean of triplicate experiments and the shaded areas represent ± one standard deviation. (**C**). From A and B, we extracted the maximal growth rate (error bars represent ± one standard deviation) of the strains that were exposed or not exposed to light. See also Figure 5, which shows the same growth rate was obtained for the same strain in liquid culture.

### Supplementary Figure S10

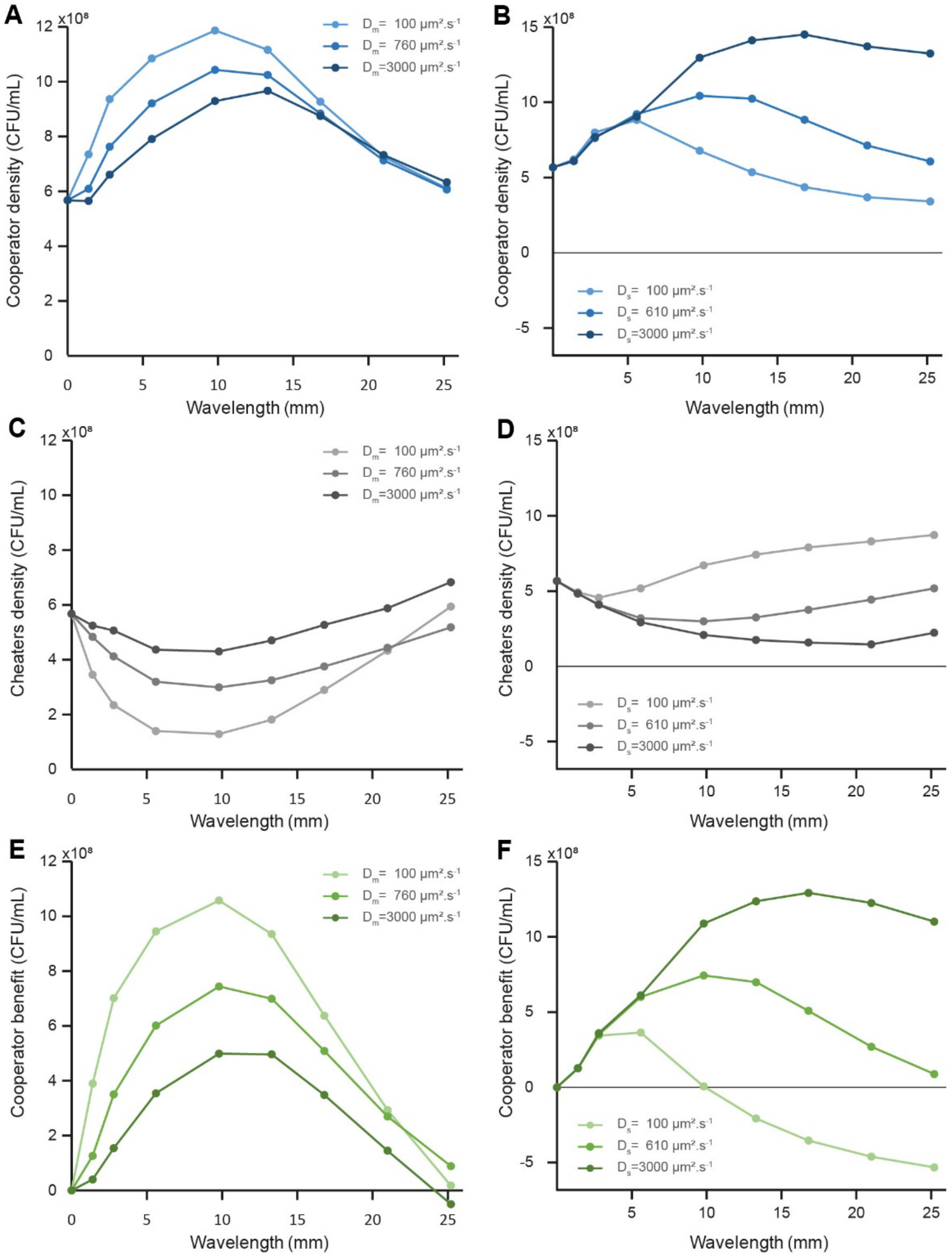

**Supplementary Figure S10**. Numerical testing of the dependency of the diffusion coefficients of glucose and sucrose on the lower and larger cut-offs of the cooperator/cheater spatial filtering properties. (**A**) The lower the diffusion of glucose, Dm, the smaller the lower cutoff $\lambda_{-}$, consistent with a shorter length-scale of diffusion of glucose, which limits cheaters' growth. The impact of the glucose diffusion coefficient on the larger cutoff was less pronounced. (**B**) The lower the diffusion coefficient of sucrose, Ds, the smaller the larger cut-off $\lambda_{+}$. This is consistent with sucrose diffusion setting the length-scale of competition within the cooperator domain. The sucrose diffusion coefficient had limited (if any) impact on the lower cutoff.

### Supplementary Movie SM1

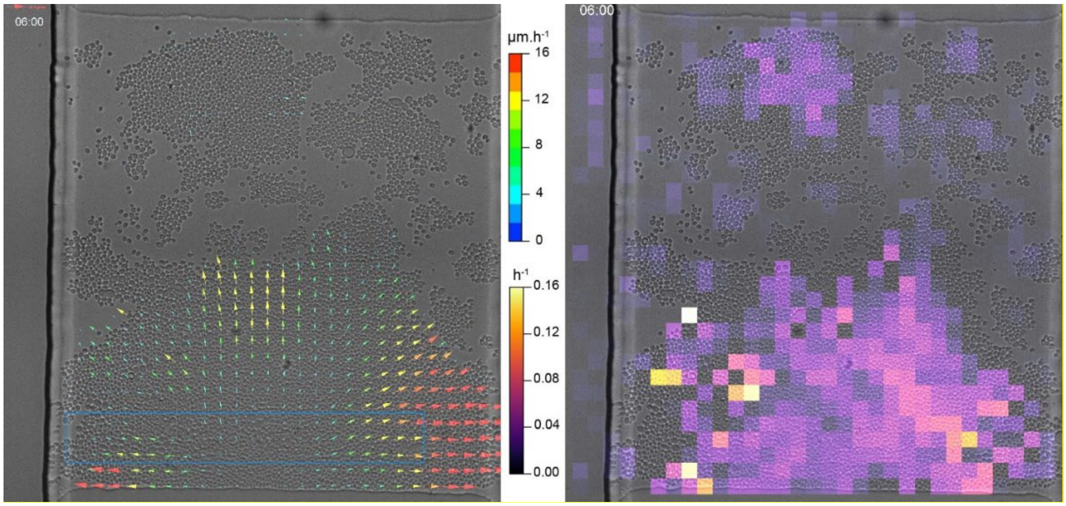

**Supplementary Movie SM1. Spatial control of yeast growth in a microfluidic chamber.** OptoSUC2 yeast cells grow in chambers of 400 µm² and are perfused via two large channels on the sides of the chambers. The height of the chamber (3.5 µm) ensures that cells grow as a monolayer. A digital micromirror device is used to project a pattern of light in the field of view (blue rectangle illuminated at 460 nm for 200 ms every 6 min). Time-lapse bright-field images were analyzed through PIV (particle image velocimetry) to generate a displacement vector map (left). We also computed the divergence map of the vector field, which is a proxy for local cell growth (right).

### Supplementary Movie SM2

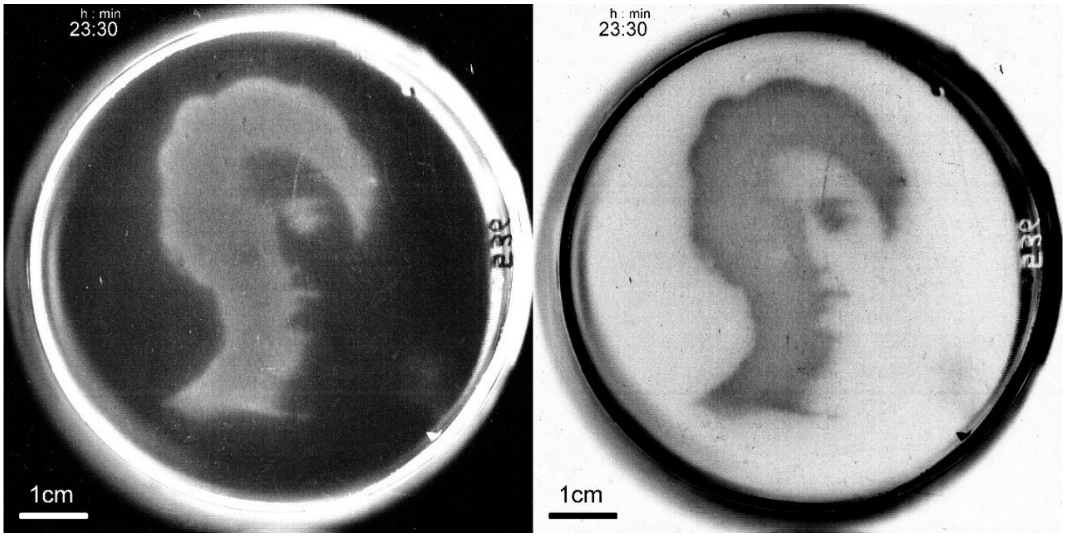

**Supplementary Movie SM2. Reproducing a picture with optogenetics-enabled yeast growth.** We projected an image of Maud Menten as a tribute to her work on the Michaelis-Menten enzymatic kinetic equation using invertase as a model. Her portrait was projected from the DMD onto a Petri dish containing OptoSuc2 cells for 45 h and the plate was scanned periodically. The yeast grew in a pattern mirroring the portrait. The first image of the timelapse was subtracted as the background to obtain the movie on the left. On the right, inverted images of the resulting yeast growth reveal the developed image of Maud Menten through OptoSUC2 light-induced growth.

### Supplementary Movie SM3

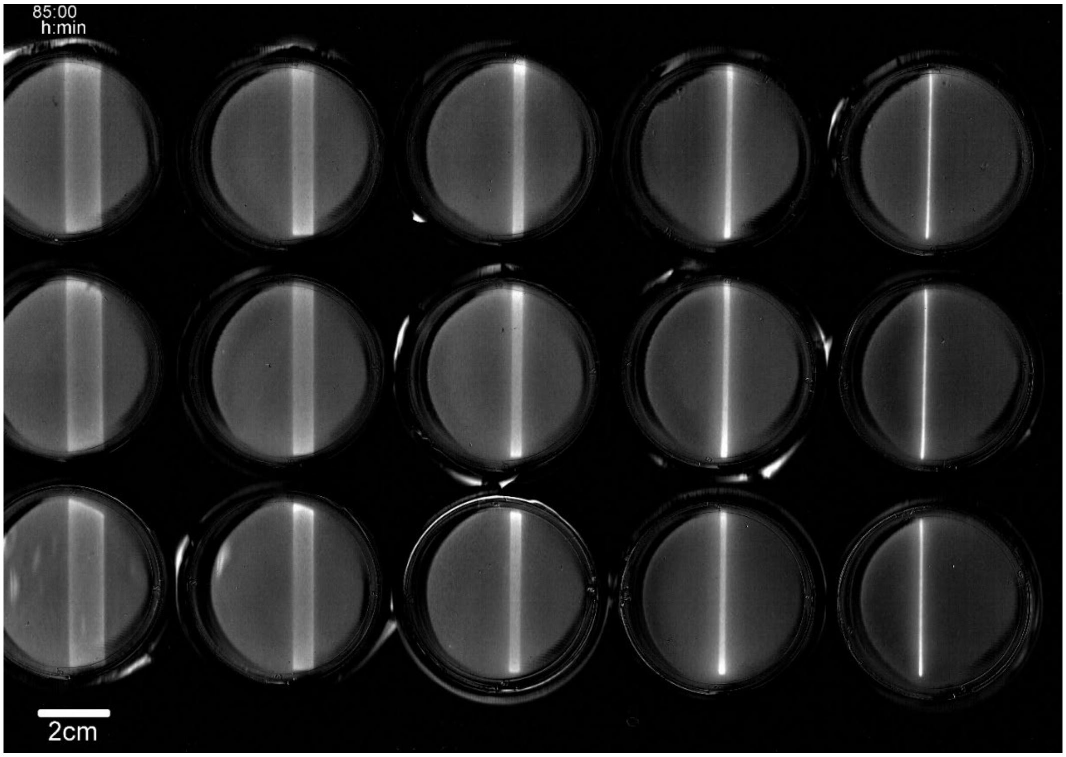

**Supplementary Movie SM3. Varying the size of the cooperator domain impacts the density of cooperators.** We projected single lines of light with varying widths and observed the growth of cells in the cooperating (illuminated) and cheating (dark) domains over time. Background subtraction was performed using the first image in the timelapse. Increasing the width of the cooperator domain both decreased the final density of cooperators at the center of the line and increased the density of cheaters at the frontier of the dark and illuminated domains. Initially, hexoses are produced everywhere in the cooperating domain, which promotes the growth of cooperating cells, decreases the sucrose concentration, and creates a source of hexose (public good). This leads to competition for glucose and an increase in the density of cheaters located at the frontiers of the cooperating domain. Within large cooperating domains, competition for sucrose leads to a decay in the sucrose concentration towards the center and an increase in the density of cooperating cells at the frontiers with the cheater (dark) domain.

### Supplementary Movie SM4

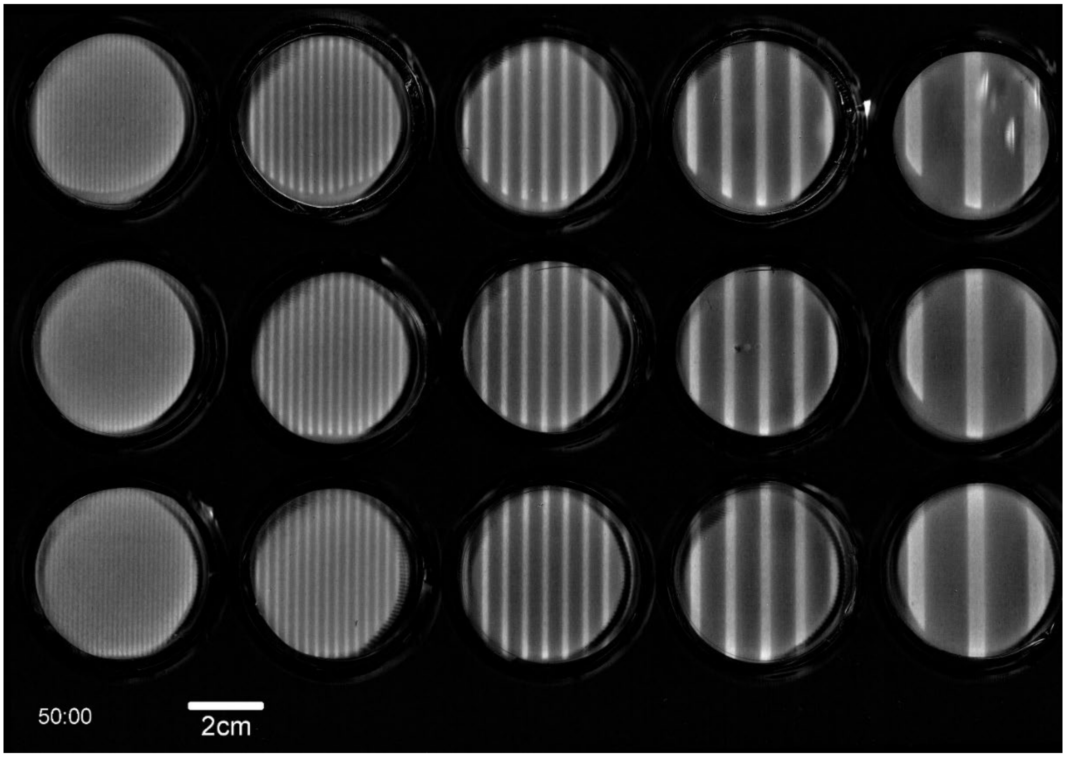

**Supplementary Movie SM4. Periodic patterning of cooperator and cheater domains gives a quantitative measurement of cooperation and competition length-scales.** We projected periodic lines of light of various widths, maintaining a constant ratio between dark and illuminated domains (75% dark, 25% illuminated). The first image was used as background measurement. Cooperators and cheaters grow in their respective domains, but the cooperator cells have a marked fitness advantage. Below the lower cutoff ($\lambda_{-}$ ~ 7 mm), the cooperator domains are too small to retain all of the glucose they produce for their own profit and the cheater cells in the dark areas can grow. Above the larger cutoff ($\lambda_{+}$ ~ 17 mm), the cooperator domain is too large to be sustainable everywhere given the limited influx of sucrose from the frontier. As a result, the cells in the central part of the cooperator domain are in competition for both sucrose and glucose (self-competition), which leads to a lower final cooperator cell density.
